# Online extinction and novelty-triggered recovery of life-long visual memories in navigating ants

**DOI:** 10.64898/2026.02.17.705905

**Authors:** Leo Clement, Cody A Freas, Antoine Wystrach

## Abstract

Insects exhibit flexible memory processes, yet current models often confine these modulations to discrete, reinforced events. During navigation, animals can learn to associate locations with resources or aversive events. However, such events occur sporadically within an otherwise continuous behavioral process involving latent learning. This raises a fundamental question for insect cognition: can insects continuously update visual memories during navigation? We show that navigating ants (*Myrmecia midas*) dynamically revalue visual memories en-route, even in the absence of any motivational change or external reinforcement. Using a trackball system that prevented visual progress, we found that nest-view attraction declined over time, reflecting a gradual, online extinction of these otherwise life-long attractive visual memories. These memory updates were view-specific, independent of path integrator state, and persisted after 24-hours, indicating the involvement of long-term memory processes. However, brief exposure to a novel visual scene was sufficient to partially restore nest-views attraction, showing that extinction is labile, and the original memories are preserved. Unlike classic paradigms, visual memory extinction operates here as a continuous, time-dependent process without reward-omission or prediction error; whereas memory reinstatement is triggered by discrete visual changes. We propose that these dynamics are mediated by opponent-processes in the insect mushroom bodies, where temporal estimates of view exposure drive extinction and novel view detection gates memory recovery.

## INTRODUCTION

Learning theory posits that associations between conditioned stimuli (CS) and biologically significant outcomes (US) are not fixed but dynamically shaped by experience. When a CS is repeatedly encountered without its expected US, such as a feeding location that no longer yields reward, the responses gradually weaken, in a process known as extinction learning. Importantly, extinction does not erase the original memory but instead forms a parallel inhibitory memory trace, allowing extinguished responses to recover following time or contextual change. These dual-memory systems enable adaptive flexibility, balancing short-term suppression with long-term retention. Across taxa, including insects, such revaluation is mediated by dopaminergic circuits signalling prediction errors.

In vertebrates, memory systems operate online during behaviour, with hippocampal spatial representations and dopaminergic prediction-error signals dynamically revaluing memories (Schultz 2016; Stachenfeld et al. 2017). In insects, studies show the formation of parallel aversive and appetitive memory traces, mediated by recurrent dopaminergic circuits in the mushroom bodies. Extinction dynamics within this known circuitry are well-established and occur when a previously reinforced stimulus is no longer presented with the reinforcer (Aso et al. 2014; Eisenhardt 2014; Felsenberg et al. 2018). However, such extinction is typically studied in a discrete, trial-based context, where the presence or absence of reinforcement events is clearly structured. Navigation presents unique challenges, where route memories guiding directed movement are formed in a continuous process through latent learning, and without explicit reinforcement (Clement et al. 2024). In contrast to associative learning paradigms, there is no clear reward or reward-omission signal during navigation between goals. We therefore asked, can such route-based memories undergo a form of online extinction during continuous, unreinforced experience?

Ants rely on sophisticated visual memory systems to establish and adjust their foraging routes, via acquiring multiple view memories which guide movement to goals (Clement et al. 2024; Mangan and Webb 2012; Schultheiss et al. 2016; Wystrach et al. 2020a; Freas et al. 2019). While navigational memories are known to be malleable, their modulation is typically studied as a discrete process tied to reinforcers or motivational changes (Freas et al. 2022; Wystrach et al. 2020a). For example, aversive learning studies show that memories can be altered by negative experiences along the route or failure to reach goals but these changes are usually assessed across multiple trips, not within them (Collett 2014; Deeti et al. 2023; Freas et al. 2022; Le Moël and Wystrach 2020; Murray et al. 2020; Schwarz et al. 2020; Wystrach et al. 2020a, 2019). “Rewinding” experiments, where ants are displaced further back along their route, causing them to experience the same views multiple times without reaching the nest, suggest within-trip memory adjustments (Collett 2014; Schwarz et al. 2020; Wystrach et al. 2019). However, such manipulations involve discrete visual changes and capture events, making it difficult to disentangle purely continuous online updating from memory changes driven by experimental interruption or consolidation during transfer.

Recent work exploring latent learning in ants has shown that visual memory acquisition in a novel environment (the initial forming of the memory) occurs continuously during route exploration even in the absence of explicit reward or punishment, yet its behavioral expression is gated during the initial acquisition phase (Clement et al. 2024). This suggests that the insect brain can support the continuous acquisition of visual memories en-route. Yet a key gap remains; latent-learning memory, by definition, is not used as it is being formed, thus its expression is only revealed during subsequent tests of route knowledge. This makes functional sense for an ant learning a novel scene, because an instantaneous recall of visual memories as they are being built would prevent the insect from distinguishing what is familiar from what is new. However, a more gradual recall, slowly increasing through time, could be viable and potentially beneficial when it comes to ignoring previous memories that turn out to be maladaptive given evidence building through time (Clement et al. 2024). What remains unclear is whether such modulation reflects cumulative online updating during navigation, or discrete changes occurring between navigational episodes.

To investigate these questions, we used a trackball system (Clement et al. 2025, 2024; Dahmen et al. 2017; Murray et al. 2020) to test if navigating bull ants (*Myrmecia midas*) can update their view memories enroute when walking in the goal direction is possible but actual visual progress is prevented. Ants could rotate freely on the trackball while on their established route, but translational movement did not result in forward displacement, preventing visual progress. This allowed us to isolate the effects of time and experience on visual memory expression in the absence of external scene changes. *M. midas* is an ideal model for this question, as it relies heavily on life-long visual route memories and often ignores alternative navigational strategies, including path integration (Freas et al. 2018, 2017a) which is typical for ants of this genus (Narendra et al. 2013; Narendra and Ramirez-Esquivel 2017).

We found that ants’ orientation toward nest-aligned views deteriorated over time, indicating an online inhibition of specific view memory attraction that overrides their long-term attractive valence. Critically, this behavioural change could not be explained by the accumulation of the path integrator. Comparisons between ants trained on open versus cluttered routes revealed that these updates were highly view-specific (Supplemental Figure 1) rather than reflecting a general reduction in motivation, implicating Kenyon cell–based representations and thus the mushroom bodies as the likely site of modification. Strikingly, these changes persisted for at least 24 h, supporting their integration into long-term memory valence rather than short-term habituation. However, transient exposure to a novel scenery, triggered a partial recovery of the original route-following behaviour when returned to the previously extinguished scene, indicating that this revaluation remains reversible within a single experience.

## RESULTS

### Orientation declines during prolonged exposure to fixed scene

Two cohorts of ants (*Myrmecia midas*) from the same nest were allowed to forage along either their naturally open route with high visual consistency (*Open* condition), or trained for 4 weeks with experimentally added visual clutter to provide distinct local visual changes towards the middle of the route, *Cluttered* condition (Figure 1A; Supplemental Figure 1).

**Figure 1.**
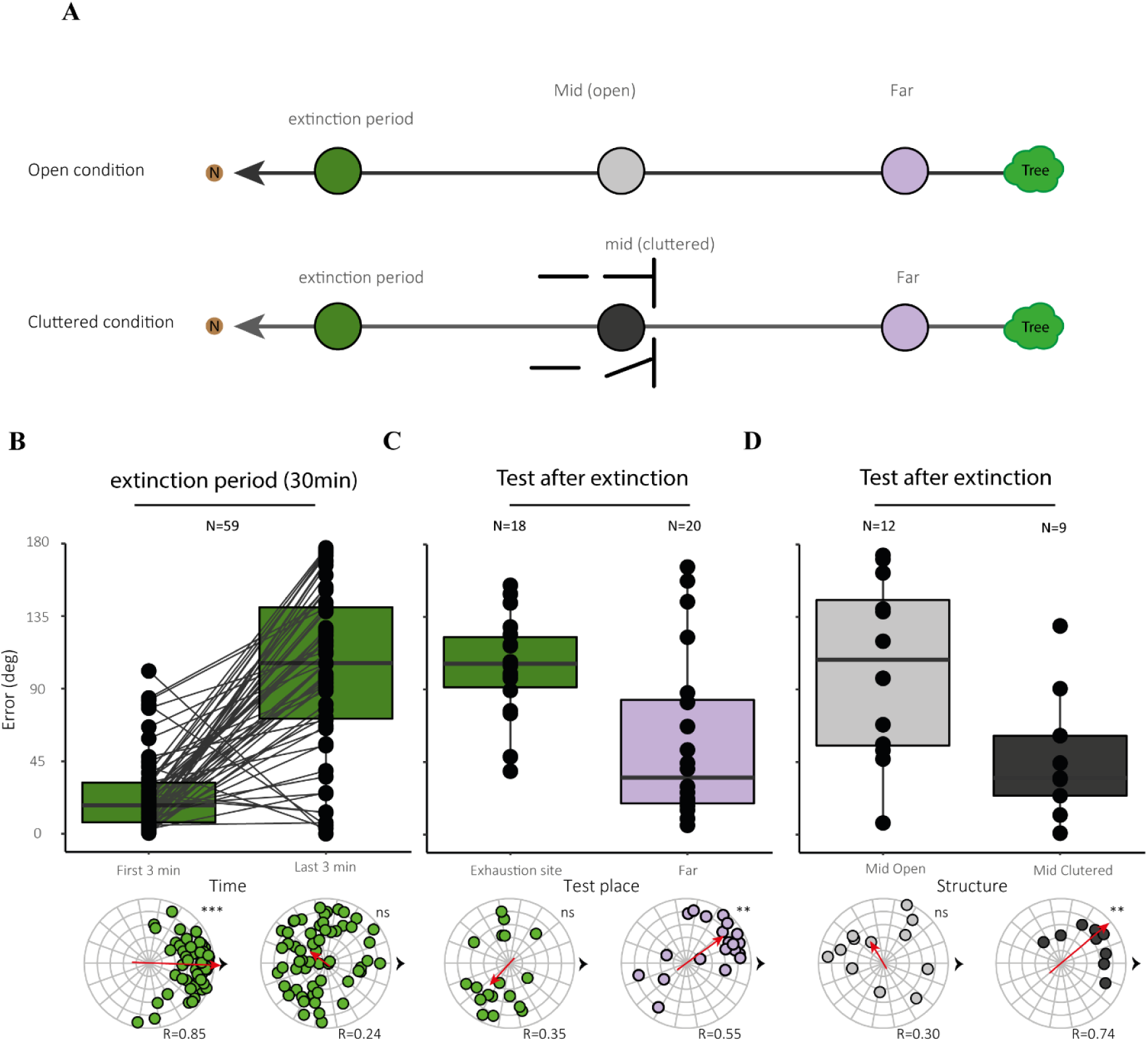
Visual extinction is context-specific and does not impair distinct route memories. (A) Experimental Procedure: Schematic of the exposure and testing protocols. Foragers were monitored for 30min at the extinction site under one of two visual conditions: an open environment or a cluttered environment. Following a 1min dark interval for transfer, ants were tested at one of three route locations: *Nest*, Mid-route (*Mid*), or Far-route (*Far*). (B–D) Orientation Dynamics. The top row shows the distribution of angular errors; bottom rows show mean directions of movement. All analyses were conducted using 3min windows. (B) Extinction Phase. Comparison of orientation at the onset (first 3 min) and the conclusion (last 3 min) of the 30min exposure. (C) Comparison of test performance for foragers recorded at the Nest versus those recorded at the *Far* site. (D) Comparison of orientation at the same test location (mid) following extinction in either an open or cluttered visual environment. Circular plots of individual angular headings for each time window and condition. Each dot represents an individual’s mean heading, with distance from the center indicating mean vector length (straighter paths lie closer to the periphery). R values (0–1) indicate the population mean vector length. Red arrows show the population mean vector direction and length; black arrows indicate the nest’s direction (to the right in all plots). Asterisks (*) denote significant orientation; ns indicates no significant orientation.

After training, homing ants were tethered on a trackball at a fixed location along their familiar nest-ward route, ∼1.5m from the nest, for 30min. While on the trackball, ants could freely rotate both their body orientation and gaze direction, experiencing rotational changes of the visual scene, but walking forward did not produce translational movement, and ants were thus prevented from making visual progress toward the nest.

As expected, ants in both *Open* and *Cluttered* conditions initially oriented toward their familiar nest-ward views when placed on the trackball (First 3min: V-test at 0°, p < 0.001, Figure 1B, dark green). Yet over the 30min tethered on the trackball, ants showed a decline in nest-ward orientation, marked by progressive increases in angular error (ANOVA: First 3min vs Last 3min: F_(1,118)_ =74.941, p < 0.001), such that by the last three minutes (27 to 30min), their headings were no longer aligned with the nest (Last 3min: V-test at 0°, p=0.980 Figure 1B). This decline of nest-ward orientation was not due to a general loss of motivation caused by the time spent walking on the trackball. Indeed, a control group showed excellent nest-ward orientation when the trackball is placed on the familiar route after spending 1hour walking on the trackball in an unfamiliar environment (see Supplemental Motivation Control). Therefore, the decline of nest-ward orientation results from the sustained exposure to the familiar visual scene in the absence of reward or navigational progress.

To disentangle whether this drop in nest-ward orientation reflects a reduction in the use of homing views in general or is specific to the views experienced during the 30min exposure period, we perform two different displacement conditions. After the 30min extinction period, ants, still tethered on the trackball, were covered in darkness for 1min while the apparatus was moved to one of three locations along their familiar route (Figure 1A): 1.) The same location near the nest where the 30min exposure happened; 2.) *Mid* site, 3m from the nest along the route; 3.) *Far* site, 6m from the nest near the foraging tree, marking the beginning of these ants’ ground-based homing route.

Ants tested at the extinction location (1.5m from the nest) remained non-nest oriented (consistent with the end of the view extinction phase; Figure 1C, dark green; V-test at 0°, p >0.05). In contrast, ants tested at the *Far* site, in both *Open* and *Cluttered* conditions, showed a strong reduction in angular error compared to the end of the extinction period (Figure 1C, purple; ANOVA *Nest* vs *Far*: F_(1,38)= 11.213,_ p = 0.002) and recovered the nest-ward orientation (V-test at 0°, p = 0.002). Thus, moving the ants into a sufficiently different visual scene along their familiar route restored visual homing as these view memories had not been part of the extinction phase.

*Mid*-site testing revealed a difference between open and cluttered routes. In the *Open* condition, where the scene in *Mid* remains relatively similar to the exposure site, ants kept high angular errors and exhibited a lack of nest-ward orientation (V-test at 0°, p > 0.05). In contrast, in the *Cluttered* condition, where the scenery in *Mid* was highly different from the extinction site (Figure. S1, *Cluttered*), ants recovered their nest-ward orientation (V-test at 0°, p = 0.007; Figure 1D, dark grey) and showed significantly lower angular error than ants in the *Open* condition (ANOVA *Mid Open* vs *Mid Cluttered*: F_(1,21)_= 6.077; p = 0.023, Figure 1D, light grey).

Path integration cannot explain these differences, as ants released in *Far* (both *Open* and *Cluttered*) or *Mid* (*Cluttered*) oriented nest-ward despite similarly large conflicting path integrator vector accumulated during the exposure period (PI length mean ± SE: *Far* =38.61±3.85m; *Mid-Open* = 38.34± 5.99 m; *Mid-Cluttered* = 53.46±6.63 m).

These results demonstrate that the declines in nest-ward homing observed during the extinction period is view-specific rather than a global reduction in homing motivation, a general exhaustion of visual-homing memories, or a conflict with the path integrator. This shows that sustained exposure to familiar views result in a progressive drop in the recall of the memories associated with these views. Ants perform a time-driven, online modulation of the expression of these memories.

#### Loss in view memory expression is long lasting

To assess the stability of the changes in memory expression following sustained exposure to a visual scene, we tested ants after a 24-h delay. Ants from a new colony were assigned to two groups. *Re-exposed* ants first experienced a 15min familiar scene exposure period on the trackball on Day 1 (1st Exposure 0h), followed by 24h in darkness, and were then placed back onto the trackball for a 3min test on Day 2 (2nd Exposure 24h). *Control* ants were collected in parallel on Day 1 but were placed directly into darkness for 24h and were tested for the first time on Day 2 (1st Exposure 24h).

*Control and Re-exposed* show a clear nest-ward headings during the first 3min on the trackball- (1st Exposure 0h: V-test at 0°, p<0.001; *Control* (1st Exposure 24h): V-test at 0°, p < 0.001). With comparable orientation during their initial exposure to the trackball (ANOVA post hoc tests, 1st Exposure 0h 0-3min vs *Control* (1st Exposure 24h), p=0.684, Figure 2A, B, green and light blue). This confirms that the 24-h delay in darkness itself did not reduce homing motivation. As expected, view exposure on Day 1 caused *Re-exposed* ants to display a gradual decline in orientation over the 15min period, with the final three minutes (12–15min) showing significantly increased angular error (ANOVA post hoc tests 1st Exposure 0h 0–3min vs last 3min, p<0.001; Figure 2A,B, light blue) and a loss of nest-ward orientation (1st Exposure 0h last 3min V-test at 0°, p>0.05).

**Figure 2.**
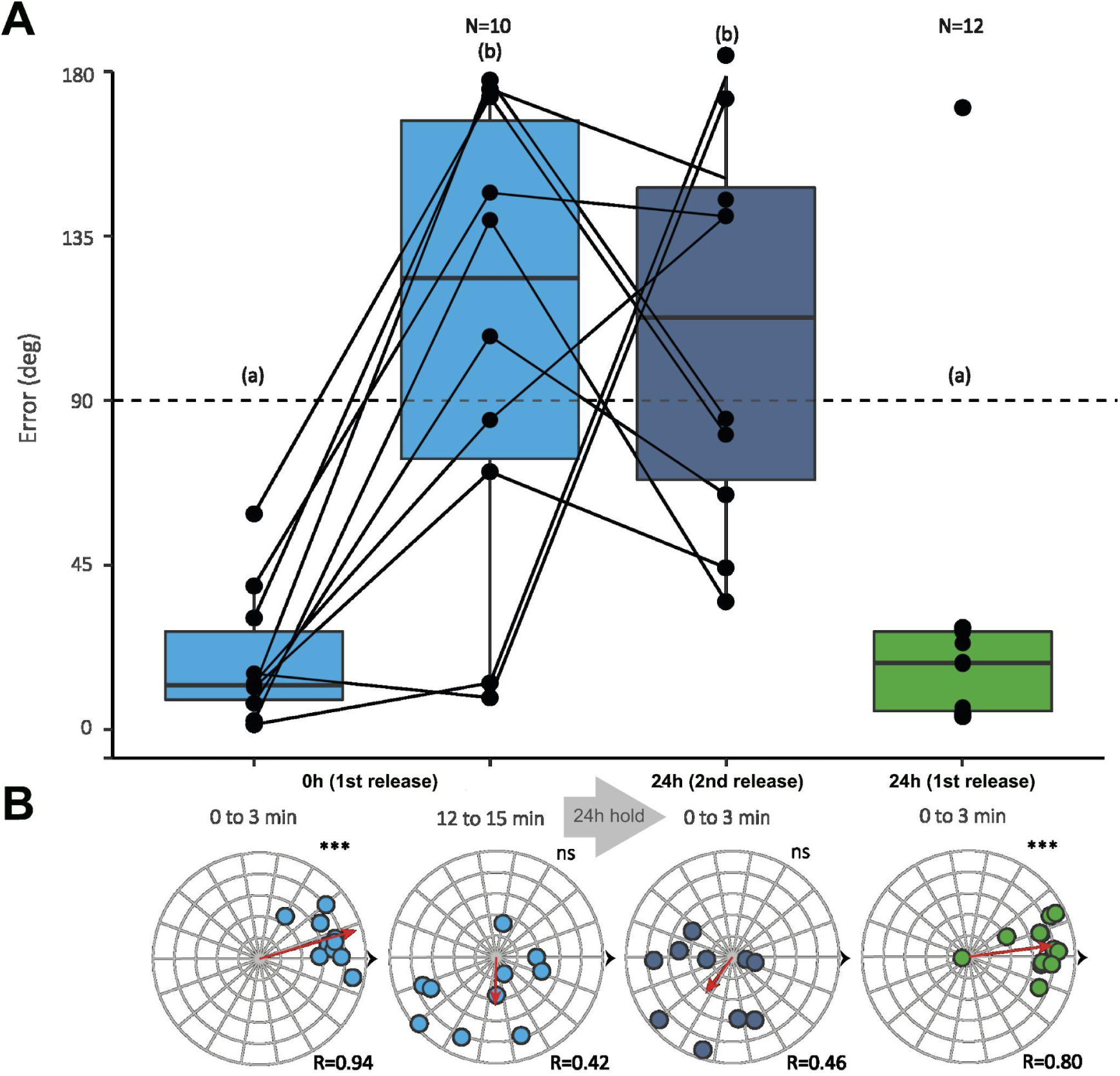
Persistence of memory valence changes following a 24-h delay. To test the stability of extinction-induced changes in visual memory valence, ants were tested 24h after initial exposure. *Re-exposed* ants were tethered on the trackball at the familiar route location on Day 1 for a 15min extinction session (0h, 1st Release), then held in darkness for 24h and tested again on Day 2 (24h 2nd Release). *Control* ants were collected at the same time but remained in darkness for 24h and were tested for the first time on Day 2 (24h 1st Release). (**A**) Distributions of angular error (top row) and mean movement direction (bottom row) for each group and time window. For *Re-exposed* ants on Day 1, we compare the first 3min (initial orientation) with the last 3min (12–15min; post-extinction). On Day 2, we compare the first 3min of *Re-exposed* ants with the first 3min of *Control* ants. (**B**) Circular plots of individual angular headings for each time window and condition. Each dot represents an individual’s mean heading, with distance from the center indicating mean vector length (straighter paths lie closer to the periphery). R values (0–1) indicate the population mean vector length. Red arrows show the population mean vector direction and length; black arrows indicate the nest direction (to the right in all plots). Asterisks (*) denote significant orientation; ns indicates no significant orientation.

Crucially, when tested again on Day 2 (2nd Exposure 24h), *Re-exposed* ants remained non-oriented to the nest (V-test at 0°, p > 0.05; Figure 2B). Their angular error was both unchanged relative to the last 3min of Day 1 and significantly higher than that of Day-2 *Control* ants (ANOVA post hoc tests 2nd Exposure 24h 0-3min vs 1st Exposure 0h last 3min: p=1.0; 2nd Exposure 24h 0–3min vs C*ontrol* (1st Exposure 24h): p < 0.001; Figure 2A, dark blue). Thus, orientation declines induced during the exposure phase are not a short-lived effect but reflect long-lasting modification of the expression of the visual memories involved.

#### Exposure to a New Scene Triggers Recovery of Visual Memory

We next investigated whether this progressive loss in memory expression reflects a transient extinction where the original memory trace remains and can be reinstated, or a forgetting process where the original memory is being erased or overridden for good. The 24-h re-exposure experiment showed that withholding visual input for a full day between sustain exposure period and test did not restore the expression of the view memories. If the original memory trace is preserved, what kind of event could reinstate its expression? Consistent with recent findings in visual latent learning (Clement et al. 2024), we hypothesised that experiencing a discrete scene change towards an unfamiliar area may provide such a trigger. We thus conducted a final experiment in which ants experienced extended exposure at a fixed familiar site along their route, followed by a transition to a novel, unfamiliar scene and then back to the original familiar exposure site.

We recorded ants (from the same colony as experience 1) for one hour on the trackball positioned at the midpoint of their route (Nest 1, 3m from nest entrance). Identical to our previous experiments, ants initially oriented strongly toward the nest (Figure. 3A; first 3min: V-test at 0°, p < 0.001) but mean angular error climbed steadily over time, reaching random chance levels (>90°) by ∼20 min, when ants were no longer significantly oriented toward the nest (Supplemental Figure 2). This significantly increased angular error, coupled with a lack of nest-ward orientation persisted for the remainder of the one-hour recording (Figure. 3A; first 3min vs. last 3min: post-hoc tests, p < 0.001; V-test at 0°, p = 0.862). Importantly, control experiments with ants tested for the same duration at an unfamiliar site rule out reduced motivation and path-integrator conflict as causes of this effect (see Supplementary Methods & Results, Figure S1; full statistics reported in the supplement).

**Figure 3.**
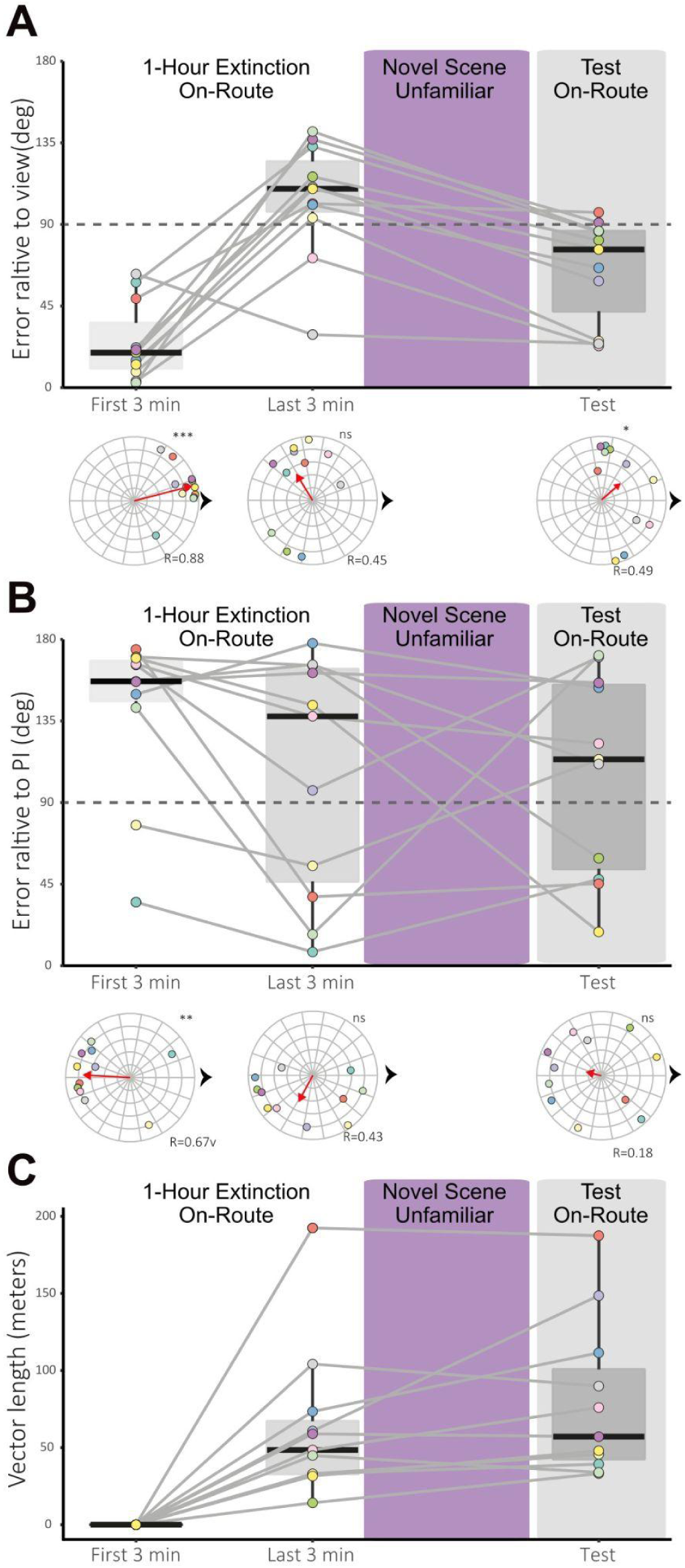
Impact of 1-hour extinction on-route on visual homing and path integration (PI) dynamics. Summary of forager headings at the start (first 3 min) and end (last 3 min) of the 1-hour extinction period at the familiar site. **(A)** Orientation vs. Nest View. Box-and-whisker plots of heading error relative to the nest-aligned direction. Foragers were initially well-oriented but showed significant deterioration in accuracy after 1 hour of extinction. Nest alignment partially recovered following a 5-minute exposure to an unfamiliar scene (purple) and subsequent return to the familiar site (grey). **(B)** Orientation vs. PI Direction. Heading error relative to the accumulated PI vector. Despite the accumulation of long vectors (see panel C), foragers showed no group-level orientation toward the PI direction at the end of extinction, during the novel scene exposure (purple), or during the final test (grey). **(C)** PI Vector Length. Distribution of accumulated vector lengths at beginning and end of 1-hour exposure period and during test. PI distance remained high and did not decrease during the unfamiliar scene exposure (purple box), indicating that the recovery of nest-aligned orientation in (A) is driven by visual memory reevaluation rather than “running down” a conflicting PI. In all plots, boxes represent the interquartile range (IQR), horizontal black lines indicate medians, and whiskers extend to the min and max. Circular plots of individual (each dot) angular heading for each time segment. The dots show both the mean direction of the heading and each individual’s mean vector length (i.e., a point closer to the periphery indicates straighter paths). The R values (range 0 and 1) indicate the population level length of the mean vector. The red arrow in each circular plot represents the mean vector direction and length of the population. Black arrow, nest direction. * Represents significant orientation while ns represents a lack of orientation. For 1min time intervals at Familiar and Unfamiliar, see Figure S2, S3.

After this one-hour view exposure period, tethered ants were covered and transported in darkness (while remaining on the covered trackball setup) to an unfamiliar site (Site A, SFigure 1). There, the cover was removed, and individuals were let to navigate on the trackball in the novel unfamiliar visual scene for five minutes before being re-covered and returned to the familiar route location for testing. Here, ants showed some evidence of the expression of the original long-term attractive memory of their familiar route: their angular error relative to the nest decreased significantly relative to the end of the exposure period (last 3min vs. test, ANOVA: F_(2,26)_ = 26.111, p < 0.001; post-hoc: p > 0.05), although it remained more elevated than their initial baseline before the sustained exposure (First 3min vs. test: post-hoc p = 0.005). Overall nest-ward orientation was recovered, although this was just below significance (Figure 3, V-test at 0°, p = 0.042). The partial recovery of nest-ward orientations cannot be attributed to the influence of the accumulated path integrator (see Supplemental Figure 2B,C 1h familiar), nor can the absence of a full recovery be attributed to a general drop of motivation (see supplemental Figure 3 1h unfamiliar, Motivation Control). This orientation pattern shows that the original memory trace was not erased, and can be, at least partially, reinstated. Therefore, the process at play corresponds to a form of extinction of route memories, whose expression decays steadily with exposure time but can be reinstated after a transient exposure to unfamiliar scenes.

## Discussion

While navigating home, *Myrmecia midas* ants prevented from making translational progress along their familiar route, and thus exposed to the same visual scene for an extended period, showed a gradual deterioration in their initially accurate, nest-aligned headings. This progressive loss of nest-ward orientation indicates an ongoing modification of the visual memory expression used for homing. Importantly, this effect was view-specific rather than reflecting a general decline in homing motivation or a global disruption of route memory use (Figure S2, S3), and it persisted for at least 24 h. Yet the change remained labile, as a brief exposure to a novel visual scene partially reinstated the original scene’s expression of nest-ward attraction.

Together, these findings align with the theoretical framework of *memory extinction* in several key respects. First, the expression of a memory associated with a specific stimulus (here, the nest-ward heading evoked by a given familiar visual scene) decreased progressively as ants experienced that stimulus without reinforcement (Figure 1). Second, the effect was long lasting (at least 24 h; Figure 2) and may therefore involve long-term consolidation processes that typically require protein synthesis (Eisenhardt 2014). Third, rather than reflecting simple forgetting, the original memory expression was recoverable (Figure 3), consistent with the idea that memory extinction is mediated by the formation of a parallel, antagonistic memory trace rather than erasure of the original memory trace (de Bruijn et al. 2021).

However, our results also differ from canonical extinction paradigms in several noteworthy respects. Most extinction studies rely on Pavlovian conditioning (Dunsmoor et al. 2015), where an initially neutral conditioned stimulus (CS) becomes predictive through its association with a reinforcing unconditioned stimulus (US), and extinction is induced by repeated non-reinforced CS presentations. Here, no discrete reinforcement is involved. The visual memory guiding nest-ward orientation is not the product of explicit Pavlovian training but instead arises from latent learning (Clement et al. 2024), which operates continuously and without discrete reinforcement events. Likewise, extinction was not induced by repeated discrete trials, but emerged gradually during prolonged exposure to the same scene (Figure 1–3; Supplementary Figure 2). This implies that memory expression is being modified while it is actively controlling behaviour. Finally, recovery of the original memory, which is often spontaneous and time-dependent in classical paradigms (e.g., de Bruijn et al. 2021; Dunsmoor et al. 2015; Felsenberg et al. 2018), could occur abruptly here and was triggered by a discrete event: brief exposure to a novel scene (Figure 3). Below, we discuss these features in turn.

### Extinction and recall occur simultaneously

Most evidence for extinction comes from associative learning paradigms in which behavioural change is assessed across clearly separated learning and test phases (e.g., Bouton 2004; Dunsmoor et al. 2015). In insect navigation, extinction-like effects have been described in bees visiting no-longer-rewarded feeders, where visit frequency declines across successive trials (Eisenhardt 2014). In such cases, extinction is expressed across discrete episodes of reinforcement-based learning. Because behavioural expression is typically measured only after rest periods or after a return to the nest, it remains unclear whether memory valence is modulated online during recall, or is instead updated between trials during consolidation windows.

“Rewinding” experiments in ants, where individuals are repeatedly captured and replaced further back along their route, show a decline in route memory expression even with short inter-trial intervals (Collett 2014; Schwarz et al. 2020; Wystrach et al. 2019). However, these manipulations involve abrupt scene changes and repeated handling, which could themselves trigger learning or retrieval processes. This makes it difficult to distinguish continuous, online modulation from trial-like resets imposed by the experimental procedure.

In the present study, across all three experiments, the decline in nest-ward orientation emerged gradually while ants remained in the same visual scene and continued to express the associated visual memory (Figure S2A). This online deterioration demonstrates that memory change can occur concurrently with recall. Even if extinction depends on the formation of a parallel inhibitory trace, its behavioural impact must emerge as that trace is being built.

This resembles observations in *Drosophila* larvae aversive conditioning, where avoidance can be expressed while learning is still ongoing (Schleyer et al. 2011). In contrast to appetitive conditioning, the aversive US must remain present for behavioural expression (Schleyer et al. 2011). Our findings show that similarly “online” dynamics apply to extinction-like processes in a navigational context. A key open question concerns when and how consolidation occurs, given that extinction effects persisted for at least 24 h.

### Extinction without US omission or prediction error

A common mechanistic account of extinction is that after Pavlovian conditioning, presentation of the CS in the absence of the expected US generates a prediction error, which drives formation of the extinction antagonist memory trace (Dunsmoor et al. 2015). Consistent with this, extinction in Drosophila has been shown to involve formation of a parallel memory when the CS is presented without the US, potentially driven by prediction-error signalling (Eschbach et al. 2020; Felsenberg et al. 2018).

In insects, parallel associative memories of different valence and timescales can be formed in the mushroom bodies through dopaminergic modulation (conveying the US), producing bidirectional plasticity at Kenyon cell (conveying the CS) to MB output neuron (conveying the CR) synapses (e.g., Aso et al. 2014; Cognigni et al. 2018; Handler et al. 2019). Complex feedback circuits between these actors can support the formation of prediction errors (Eschbach et al. 2020; Terao and Mizunami 2017; Webb 2024).

Although much of this framework comes from olfactory conditioning in *Drosophila*, a similar logic in the mushroom body applies to visual learning in the context of insect navigation (Ardin et al. 2016; Buehlmann et al. 2020; Kamhi et al. 2020; Webb and Wystrach 2016; Wystrach et al. 2020b). Notably, the memory dynamics of extinction may not be identical across insect species. For instance, olfactory memories in ants show remarkable resistance to extinction protocols (Piqueret et al. 2019) whereas our results suggest that visual memories are highly susceptible to time-driven extinction. Such a divergence may reflect the distinct ecological demands of each modality, where the functional necessity for stable olfactory cues contrasts with a visual navigation system that must remain highly plastic to filter out outdated or inaccurate information.

However, the extinction observed here cannot be explained by prediction error. The original attractive route memory was acquired through latent learning, where views along the route are learned continuously without the need for explicit reinforcement such as reaching the nest or finding food (Clement et al. 2024). Thus, the visual scenes had always been experienced *without* reinforcement, so that extinction cannot be explained as a prediction-error response to US omission.

This creates an apparent paradox: if the original memory forms without a US, and extinction also develops without a US, what determines whether exposure strengthens memory expression or suppresses it?

A potential resolution follows directly from mechanistic accounts of latent learning. Clément et al. (2024) proposed that continuous route learning involves both the ongoing appetitive learning of views via continuous dopaminergic input and in parallel, continuous formation of a transient antagonistic trace that suppresses expression of the appetitive memory, potentially via lateral inhibition between MBONs. Such a mechanism would prevent premature recall during initial learning, allowing the insect to distinguish novel from familiar scenes. Retrieval of the latent memory is then triggered by discrete events such as entering the nest or experiencing a sudden change in scenery, which reset the antagonistic trace and unmask the attractive output.

Notably, this framework can also account for our extinction results (Figure 4). When ants are exposed to a familiar route scene, the attractive memory is initially expressed as nest-ward orientation (Figure 4A). Due to continuous learning, prolonged exposure to the same scene without progress makes the antagonistic trace can gradually rebuild and increasingly suppress expression of the (already learnt) attractive output, yielding the observed decline in nest-ward attraction (Figure 4B). This produces extinction-like dynamics without requiring reinforcement or prediction error which are key components of classical paradigms.

**Figure 4.**
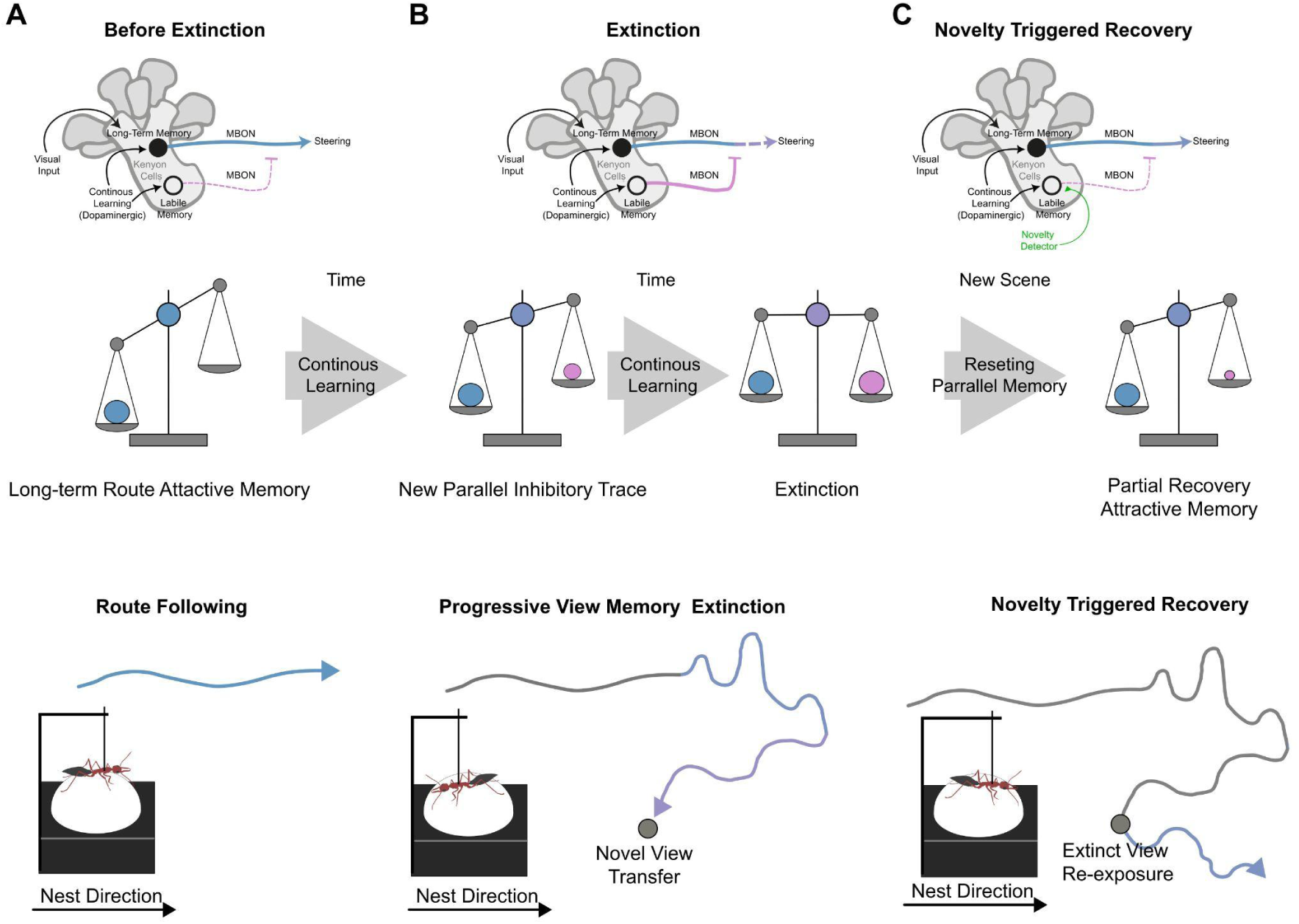
A simple Mushroom Body (MB) circuitry model captures the dynamics of time-dependent view memory extinction and context triggered recovery. The top row illustrates the proposed neural circuitry of the observed memory dynamics. The middle row depicts the relative weighting of the original long-term attractive memory, the parallel inhibitory trace that develops during prolonged exposure and the novelty-triggered recovery. The bottom row illustrates the behavioural output in the ant’s path. (**A**) Before extinction, the familiar route memory’s attractive valence dominates, producing strong nest-aligned homing. (**B**) during prolonged view exposure on the trackball, an inhibitory parallel memory trace builds, reducing the overall attractive valence of the associated view memory, leading to reduced nest alignment. As the two memory traces reach similar strength, the net attractive valence of the view declines, and nest-aligned homing deteriorates. **(C**) Via a novelty detector, the inhibitory memory trace is reset, leading to recovery of the overall long-term attractive memory trace, which leads to a partial recovery of nest-aligned homing.

### Sudden, novelty-triggered recovery of an extinguished memory

Extinguished responses often show spontaneous recovery over time, consistent with the idea that the parallel extinction memory trace is transient (Bouton et al. 2006; Rescorla 2004). In *Drosophila*, recovery can take days (Hirano et al. 2016). Here, we found that the extinction memory remained stable after 24h in darkness, it therefore remains possible that additional time would produce spontaneous recovery. Over longer periods without exposure, visual memories in ants can become functionally discounted as outdated, yet this does not imply memory decay, but rather reflects a reweighting of older visual memories as less reliable relative to more recently experienced cues (Freas and Cheng 2017).

Crucially, we show that recovery can also be triggered rapidly by a discrete event: brief exposure to a novel visual scene (Figure 3). This pattern is predicted by the latent-learning framework described above. If novelty resets the antagonistic trace, then a transient scene change should remove inhibition and immediately allow the original attractive memory to re-emerge (Figure 4C).

This event-triggered recovery broadens the range of mechanisms underlying extinction-like behavioural dynamics. One possibility is that novelty-dependent modulation acts by resetting the KC→MBON synaptic input of the extinction memory trace. Under this view, darkness may fail to induce recovery simply because it provides no novel visual input.

### Partial recovery as a consequence of continuous learning

Spontaneous recovery after extinction is rarely complete in classical paradigms (Delamater and Westbrook 2014; Dunsmoor et al. 2015). Consistent with this, recovery of nest-ward orientation in our ants was only partial: headings were biased toward the nest but less accurate than at the beginning of the procedure (Figure 3). This reduction was specific to ants that experienced the extinction procedure and was not simply due to time spent on the trackball (Supplementary Figure 2, 3).

Partial recovery could reflect residual inhibitory influence or a modification of the original memory trace (or both). The latent-learning model predicts the latter. As ants progressively lost nest-ward attraction, they explored alternative headings (Figure 3, 4B). If continuous learning remained active, then views associated with these novel directions would also be incorporated into the attractive memory (alongside formation of an antagonistic trace). When novelty subsequently resets inhibition, the expressed attractive output would then reflect a mixture of the original nest-ward direction and the additional directions sampled during extinction, yielding a weaker and less precise nest-ward bias. Overall, the model proposed for latent learning and event-triggered recall (Clement et al. 2024) can account for both extinction and recovery in our data without additional assumptions. These behavioural results therefore provide converging support for that framework.

### Inhibition versus aversion

A dominant view of extinction is that the CS acquires inhibitory properties that suppress the conditioned response, rather than becoming aversive (e.g., Bouton et al. 2006; Larrauri and Schmajuk 2008). In insect mushroom bodies, different compartments are often described as supporting appetitive or aversive memories that compete for behavioural control, although their interactions can produce more complex forms of regulation (e.g., Aso and Rubin 2016; Cognigni et al. 2018; Krashes et al. 2009; Perisse et al. 2016). In *Drosophila*, extinction of aversive memories can involve formation of competing appetitive traces, although extinction does not actually result in attraction (Felsenberg et al. 2018). This raises the question of whether the decline in nest-ward orientation observed here reflects the gradual build-up of an aversive memory opposing attraction, or merely an inhibitory trace that gates attraction without generating repulsion.

Aversive learning in ant navigation can be induced by trapping, which leads to active turns and avoidance of the capture zone (Freas et al. 2022; Lionetti et al. 2024; Wystrach et al. 2019). The resulting opponent attraction–repulsion processes have been proposed to support detours, nest pinpointing, and other navigational behaviours (e.g., Le Moël and Wystrach, 2020; Murray et al., 2020; Schwarz et al., 2020).

Our results suggest that the extinction observed here reflects inhibition rather than aversion. As ants were walking on a trackball with no change in visual scenery, an aversive memory “winning” over attraction should produce strong turning or avoidance-like behaviour. We did not observe such avoidance, and several ants randomly resampled the nest-ward orientation after having expressed extinction (Supplemental Figure 2A; Supplemental Figure 4).

Therefore, the extinction memory appears to suppress attraction without driving repulsion to these views. Functionally, this distinction is important as suppression allows ants to disengage from a failing strategy without triggering maladaptive flight responses in a familiar environment.

### Functional significance

These experiments reveal an extinction-like process in the ethological and operant context of navigation, providing insight into both mechanism and function. Ants learn their foraging routes through continuous visual learning, enabling robust shuttling between nest and foraging sites (Amin et al. 2025; Clement et al. 2024; Freas and Cheng 2025; Mangan and Webb 2012; Wystrach et al. 2013). Yet our results show that prolonged exposure to a familiar scene gradually reduces reliance on that memory and promotes exploration of alternative headings. Such extinction protects ants from becoming trapped in situations where route following is ineffective, even in the absence of overtly punitive signals: for instance, if an obstacle prevents forward progression or in cases where maladaptive looping trajectories are formed (Clement et al. 2025). In such conditions, rigid persistence on the ant’s life-long appetitive route memories could be fatal, whereas extinction-like suppression would release behavioural control to alternative strategies such as systematic search and path integration. If successful, these new directions would then be incorporated into the route memory through continued learning, explaining why, on the next trial, the ‘recovery’ is only partial and differs from the original memory expression.

## Conclusions

We describe an extinction-like process in the context of natural navigation, where prolonged exposure to a familiar visual scene without progress steadily suppresses nest-ward attraction. This extinction does not rely on reward omission or prediction error, but instead fits naturally within the dynamics of latent learning: visual memories can be continuously updated while being expressed, and rapidly re-expressed after discrete novelty-driven reset events. Functionally, this mechanism provides an adaptive way to disengage from an ineffective homing strategy and promote exploration. More broadly, our results highlight that “extinction” is not a single canonical process but can be implemented through multiple mechanisms and timescales, and that the mushroom body circuitry may provide the flexibility needed for such task-specific fine-tuned solutions to evolve.

## Materials and Methods

### Study Species

*Myrmecia midas* is a nocturnal bull ant that navigates via terrestrial visual cues along established routes between the nest (typically at the base of a tree; Deeti et al. 2024) and foraging trees. While they possess a celestial compass, panorama view memories are the primary navigational cue (Freas et al. 2017a, 2017b; Freas and Cheng 2018).

### Study site and collection

Experiments were conducted from February to May 2024 on two *Myrmecia midas* nests on the Macquarie University Campus in Sydney, Australia (33°46 11S, 151°06 40E). For all experiments, inbound foragers were collected in the pre–morning twilight as they descended their foraging tree and individually held with sugar water within phials until testing. At Nest 1 all foragers were collected from a foraging tree 5.8m from the nest and at Nest 2 all foragers were collected from a foraging tree 7.2m from the nest entrance. Testing began each day at sunrise and ran until 14:00h. Tested ants were marked with paint (POSCA^TM^) and released back to the nest.

### Trackball set–up

To record foragers’ paths, individual foragers were tethered upon one of two trackball devices (Dahmen et al. 2017; Murray et al. 2020). Each device consisted of a polystyrene ball levitated above an aluminium cup by constant air stream provided by a pump. These devices were similar in their construction and recording method but differed in their size; Trackball One: 5cm in diameter; Trackball Two: 10 cm in diameter (Murray et al. 2020). Both Trackballs were used during the one-hour extinction test (randomly assigned to individuals across both *familiar* and *unfamiliar* conditions), while only Trackball Two, which is permanently housed at Macquarie University, was used to conduct the open/cluttered and re-exposure extinction experiments.

Each device had two sensors placed at 90° to the azimuth of their spheres, which allowed the recording of the ball’s movements at 30Hz and allowed translation of this motion into X and Y data to reproduce the forager’s path. Horizontal movement of the ball was restricted by two small wheels. To tether a forager into place atop the polystyrene ball, magnetic paint was applied to the thorax of each ant. The tethering apparatus which held the ant in place consisted of a 3mm magnet which was attached to 5mm of dental thread. This thread was attached to a 0.5mm diameter carbon pin housed within a glass capillary (Clement et al. 2025; Clément et al. 2023; Dahmen et al. 2017). This allowed tethered ants to control both their gaze direction and rotate their body orientation upon the ball, with forward or backwards movement being translated to the ball’s rotation. Importantly this means that while an ant is tethered within the system, translational movement results in no visual feedback or ‘progress’ in reaching the goal location.

### Experimental Designs

#### Open and Cluttered route experiment

To test online visual memory modulation and its view specificity, we examined whether the extinction of specific views at one location influenced visual memory valence along the entire foraging route. Open, uncluttered routes maintain high visual similarity across large distances, whereas cluttered routes contain prominent local objects that make views along the route visually distinct (Zeil 2012; Zeil et al. 2003). This allows us to assess whether memory extinction is view-specific or represents generalized reductions in homing motivation across the route.

At Nest 1, the route was naturally clear of local visual clutter (Supplemental Figure 1). In this open environment, foragers were placed upon the track ball 1.5m from the nest and their paths were recorded for 30min. After 30min, a cover was placed over the ant, blocking all visual cues for 1min, and the trackball was randomly assigned and transported to one of three test locations along the foraging route: 1.) *Nest,* the same location 1.5m from the nest, 2.) *Mid*, the middle of the foraging route 3m from the nest, 3.) *Far*, the furthest point (5.8m) from the nest at the base of the foraging tree. Once the trackball was placed at one of these locations, the cover was removed, and paths were collected for 5min.

After the *Open* condition was complete, visual clutter was added to the foraging route. Wooden boards were erected along the route, focused on creating visual contrast between the *Nest* and *Mid* sites (Supplemental Figure 1). As this represented a large visual change to the route, which *M. midas* is highly sensitive to (Islam et al. 2020), foragers were allowed to become familiar with the new visual route for 28 days. After familiarisation, a separate group of ants were tested on this cluttered route with an identical methodology to open route testing. Foragers were placed upon the trackball 1.5m from the nest for 30min, covered, then tested at one of the three test locations (*Nest, Mid* or *Far)*.

#### 24-hour re-exposure experiment

To test the stability of these view memory changes, view extinct foragers were tethered to the same site 24h after extinction. At Nest 2, collected individuals were randomly assigned to a *Re-exposure* or *Control* condition. The morning of collection, *Re-exposure* assigned foragers were placed on the trackball along their foraging route ∼1.0m from the nest entrance (Supplemental Figure 1) and their paths were recorded for 15min. After this period, individuals were immediately returned to their vial and held in darkness for 24-hour, when they were recorded for a second time upon the trackball at the same location for a further 15min. To ensure nest-ward motivation remained stable after the 24-hour delay, *Control* foragers were not tested the first morning after collection. Instead, each individual was held in darkness until the next day and tested (15min) along with the *Re-exposed foragers*’ second test on the trackball.

#### 24-hour re-exposure experiment

To test whether view memory extinction persists over 24 hours, indicating long term changes in memory valence rather than short-term habituation. At Nest 2, foragers were randomly assigned to one of two groups: *Re-exposure* or *Control*. On the morning of collection, *Re-exposure* individuals were tethered to the trackball ∼1.0m from the nest entrance (Supplemental Figure 1) and their paths recorded for 15min (initial extinction phase). Following this, ants were held in darkness for 24 h. On the next day, they were placed back on the trackball at the same location for a second 15min recording. *Control* group individuals were captured and held in darkness for 24h without initial trackball exposure. They were then tested for the first time on the trackball alongside the Re-exposure group’s second test, to ensure motivation and orientation performance were comparable.

#### one-hour view exposure

At Nest 1, collected individuals were randomly assigned to one of two conditions: *familiar* and *unfamiliar (for Unfamiliar testing see Supplemental Materials)*. Individuals in the *familiar* condition were tethered upon the trackball at the mid-point of their foraging route, ∼3m from the nest and their foraging tree. Tethered individuals were allowed to travel in a chosen direction, with their paths collected for 60min. After this period, a cover was placed over the ant which blocked all visual cues. While covered, the trackball was transported to a distant, unfamiliar location (Unfamiliar Site B; Supplemental Figure 1) where the tethered ant was uncovered, and their path was collected at this site for 5min before being re-covered for another 5min while the trackball was transported back to the original familiar location along the route. Once the trackball was placed back at the centre of the foraging route and the cover removed, then the *test* began with individual paths collected for 5min at this site.

#### Data extraction and analysis

All statistical analyses were run using the open-source software R (v 3.6.2. R Core Development Team). For all statistical tests, the p-values were compared to the critical alpha risk at 0.05, with the appropriate correction if needed. The statistical parameter’s mean and the associated standard error is given within the text and/or in figures: N represents the number of individuals.

As we used two trackball configuration recording at different frequency rates, we down sample path on one of them to match 60 hz. We processed the path by first smoothing the x, y coordinate using a Savitzky-Golay filter (from R “trajr” package) with a window length of 1 seconds. Then, we extracted the angular velocity time series, which was smoothed twice: first with a moving median, then with a moving mean, both with a window length of 0.2s). All parameter estimates from our analysis were obtained by segmenting our data path into non-overlapping windows of either 180 or 10 seconds.

#### Analysis of the ants’ direction of movement

To determine extinction learning of specific views, our focus was on the direction of movement, which is a direct measurement of the ants’ ability to use view to direct her path. Within each segmented window we determine the mean direction of movement (mu) as well as the mean circular vector length (r, a measure of dispersion) of each individual. The mean directions (mu) were analyzed using a Rayleigh test (from R package: Circular) that also includes a theoretical direction (analogous to the V-test). To test whether the angular data are distributed uniformly as a null hypothesis or if they are oriented toward the theoretical direction of the nest as indicated by the familiar panorama.

As a direct reading of extinction learning, we estimated a path directional error, defined as the absolute angular error between the path orientation and the nest direction. These path directional errors were compared to the 90° threshold (which corresponds to a random direction taken by the population) using a Wilcoxon-Mann-Whitney test.

#### Analysis of the ants’ path integrators effect

To control for the effect of the path integrator within the window, we extracted the central value of the X path. This value represents the relative position of the path integrator from the ant’s perspective. A negative value indicates that the ant has followed its route and accumulated a negative vector pointing away from the nest. Conversely, a positive value indicates that the ant is heading away from the nest, thus accumulating a vector pointing towards the nest. If these reductions in nest-ward headings were due to an accumulation of a negative vector, we should expect that long negative vector should direct ants’ path to the opposite direction of the nest. To test for this effect, we used correlation between error and X path integration value.

#### Statistical models

For each experiment, we tested: the effect of the time spent on the path directional error in linear mixed model. As our individuals were recorded across several conditions (across time during extinction and test), the models are mixed models that controlled the individuality effect as a random variable. It should be noted here that in case of deviation of the residuals of these models from normality and or homoscedasticity, the response variables have been transformed. Model were subsequently analyzed via an analysis of variance (Anova), followed by a post-hoc analysis of Tukey’s rank comparison.

## Acknowledgements

We are grateful to Macquarie University for permission to work on site and access to the nests.

## Data Availability

Raw Data and Code will be made available on github: https://github.com/ClementLe0/.

## Ethics Statement

No institutional or governmental ethical approval is required for research involving insects in Australia. All procedures were strictly non-invasive and individuals were returned to their nest after testing.

