## Supplemental Methods; Supplemental Figures for "Online extinction and novelty-triggered recovery of life-long visual memories in navigating ants"

**Supplemental Motivation Control and Path Integration Analysis**

**Methods**

*One-hour unfamiliar view exposure experiment*

In the unfamiliar condition, ants were tethered upon the trackball at a distant, unfamiliar site (Unfamiliar Site A 70m from the established route; Supplemental Figure 1) with their paths collected for 60min. After 60min, a cover was placed tethered ants were covered for 5min while the trackball was transported to a distinct unfamiliar site (Unfamiliar Site B, 51 meters from the established route) where the ant was uncovered, and their path collected for 5min. Ants were then re-covered for 5min while the trackball was transported to the familiar location at the centre of their foraging route. The cover was removed, and the test began with individuals’ paths collected for 5min.

Data processing and statistical analyses

Path integration vector estimation

The state of the path integrator (PI) was estimated by summing successive displacement vectors over time to generate a home vector representing the ant’s inferred distance and direction to the nest. Vector length was normalised by total path length where appropriate. PI direction was expressed relative to the nest-aligned visual direction, such that a negative PI vector accumulated during trackball walking pointed approximately 180° from the nest direction.

To test whether PI state influenced orientation, we examined changes in PI vector length and direction across time windows and assessed whether ants oriented preferentially toward their PI vector using circular statistics.

**Results**

*Path integrator state cannot explain the increase in angular error*

We examined whether the deterioration in route-following behaviour could be explained by the accumulation of a conflicting path-integration vector. As ants walk toward the nest on the trackball at the familiar site, they accumulate a negative vector pointing backwards (180° to the nest-aligned view), which could, in principle, introduce conflict between visual and path-integrator cues. If the observed disorientation were due to the PI, ants should orient and “run off” this vector during the five minutes spent at the unfamiliar site. However, vector lengths remained stable at Site A, and foragers were never significantly oriented toward their PI vector during either the 1-h extinction period or the post-view-change test (Supplemental Figure 3). These results indicate that the increased angular error reflects modulations to visual memory rather than the interaction between views and the path integrator state.

*Unfamiliar view exposure does not reduce motivation to use visual memories*

As a control, we evaluated whether prolonged tethering alone reduces ants’ motivation to rely on visual route memories. Ants were tested for one hour at an unfamiliar site (Site A, SFig. 1), transferred to a second unfamiliar site (Site B, SFig. 1) for 5min and then transferred to the midpoint of their familiar route. During the one-hour period at the unfamiliar site, angular error did not differ from random expectations (90° threshold; Wilcoxon, p>0.05), and orientation was uniformly distributed across all time windows (Rayleigh tests, p>0.05). This confirms that Site A was genuinely unfamiliar for these foragers. In contrast, when these ants were returned to the familiar views of the route centre for the 5-min test, angular error sharply decreased, and individuals showed significant nest-ward orientation (V test; p<0.001). These test headings demonstrated that M. midas foragers remain motivated to attend to and use familiar terrestrial cues even after more than one hour on the trackball. Thus, the behavioural changes observed after extinction at the familiar site cannot be attributed to reduced motivation or fatigue but instead reflect genuine, enroute modifications to the visual memory associated with that specific view.

**Supplemental Figures**

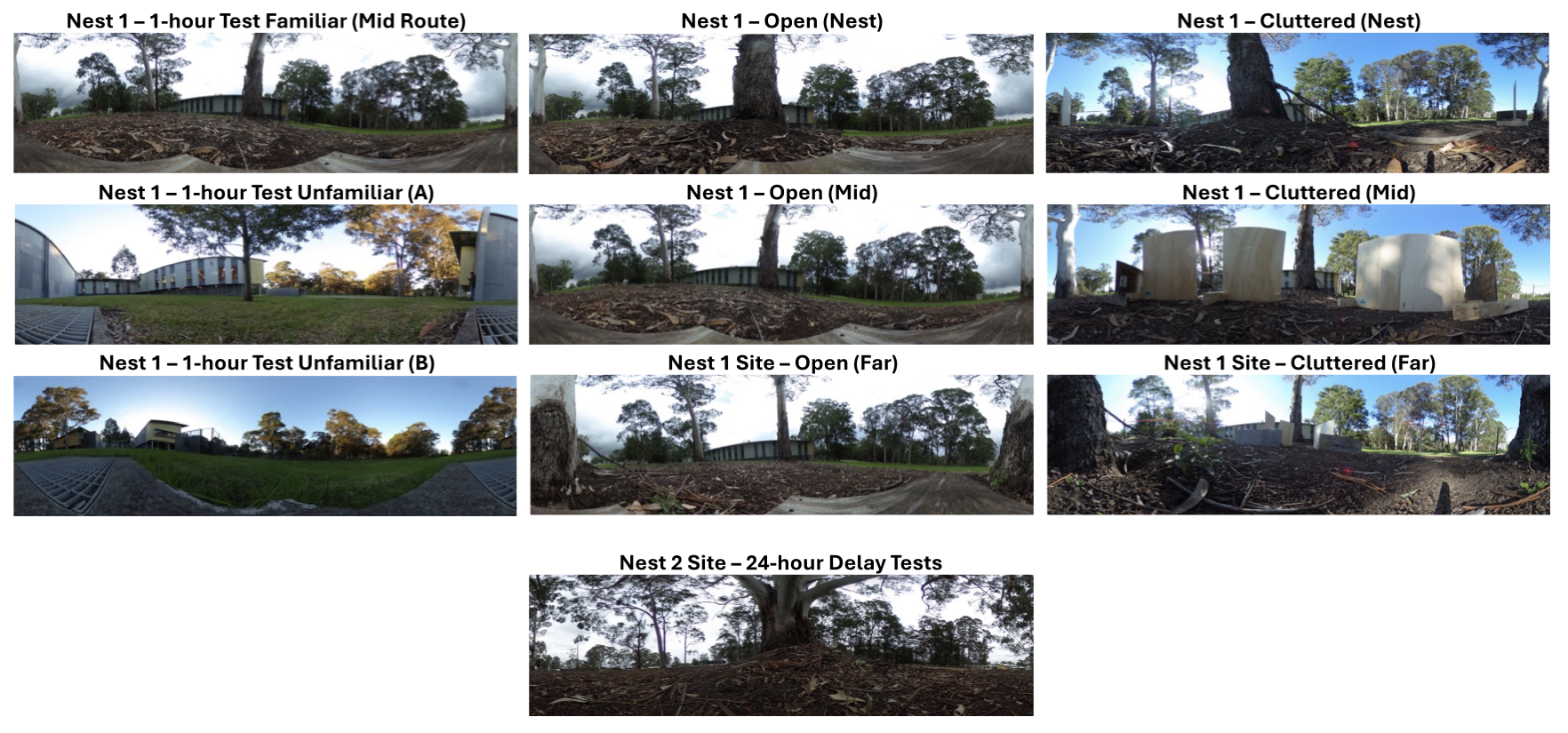

**Supplemental Figure 1.** 360º Panoramic images of all testing sites used across the three experiments. All panoramas at familiar sites are oriented to have the nest (and nest tree) centred. At the two unfamiliar sites, the panorama orientation is centred on the compass direction of the inbound route to the nest (all nest 1 panoramas are in the same compass direction).

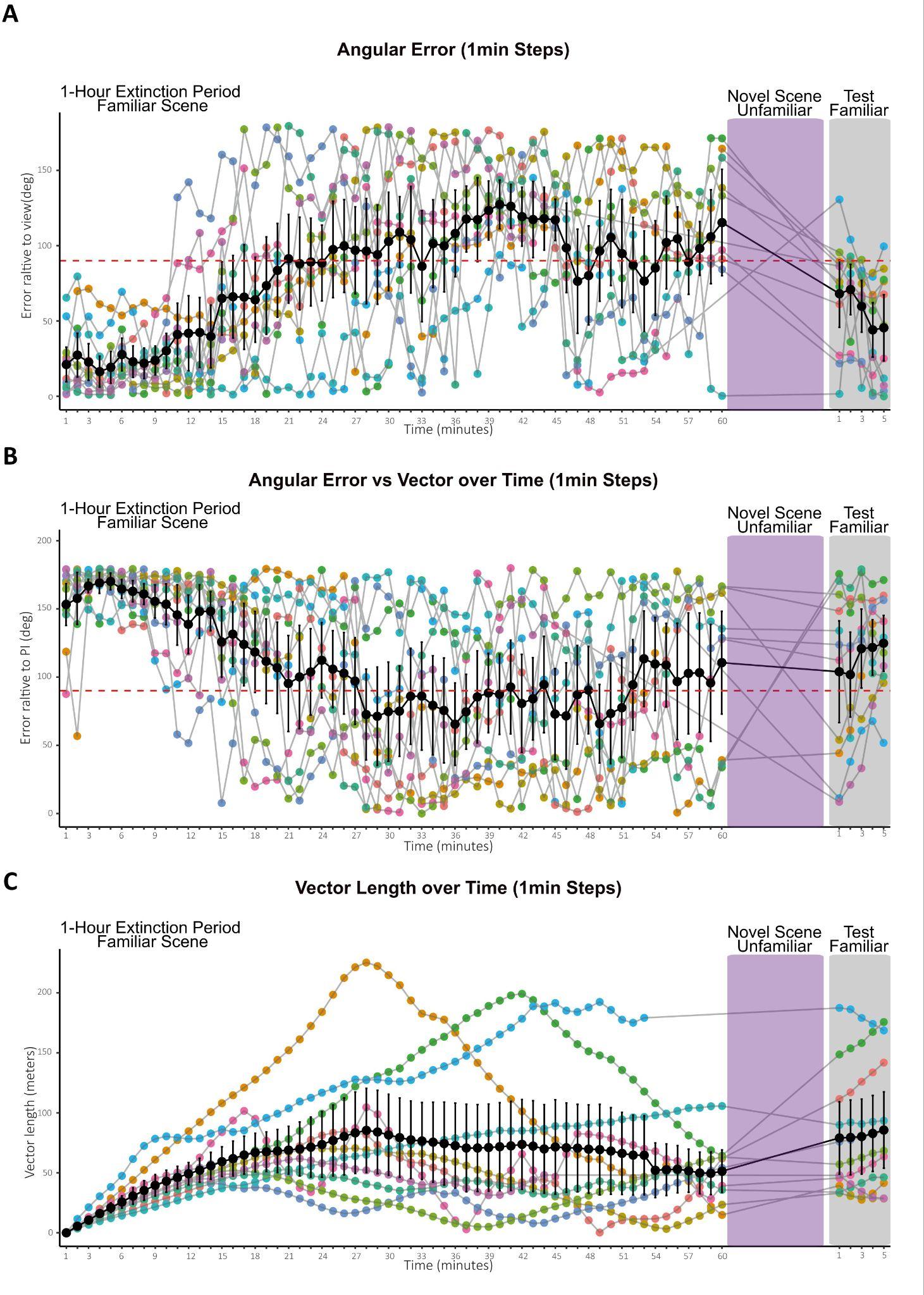

**Supplemental Figure 2. Fine-scale 1min steps in heading error relative to nest-aligned views and the path integrator (PI) over 1h extinction at the familiar site.** (**A**) Angular Error vs the nest. Foragers were initially well-oriented toward the nest-aligned direction, but orientation accuracy deteriorated over the 1-hour extinction period at the familiar site. Following a 5-minute exposure to an unfamiliar scene (purple), foragers were returned to the familiar scene for a 5-minute test (grey), resulting in a partial recovery of nest alignment. (**B**) Angular Error vs the PI direction (accumulated on trackball). Despite accumulating long PI vectors (>50m, see panel C) during the 1-hour exposure (C), foragers showed no significant group-level orientation toward the PI vector during the extinction period, the novel scene exposure (purple), or the final test (grey). (C) Vector Length. PI vector distance did not decrease during the unfamiliar scene exposure (see grey lines in purple box). This indicates that the observed recovery of nest-aligned orientation was view-memory based reevaluation rather than caused by foragers "running down" their PI memory during the 5-minute interval at the unfamiliar site. Black line, mean. Bars, SE. The red-dotted line denotes the orientation threshold.

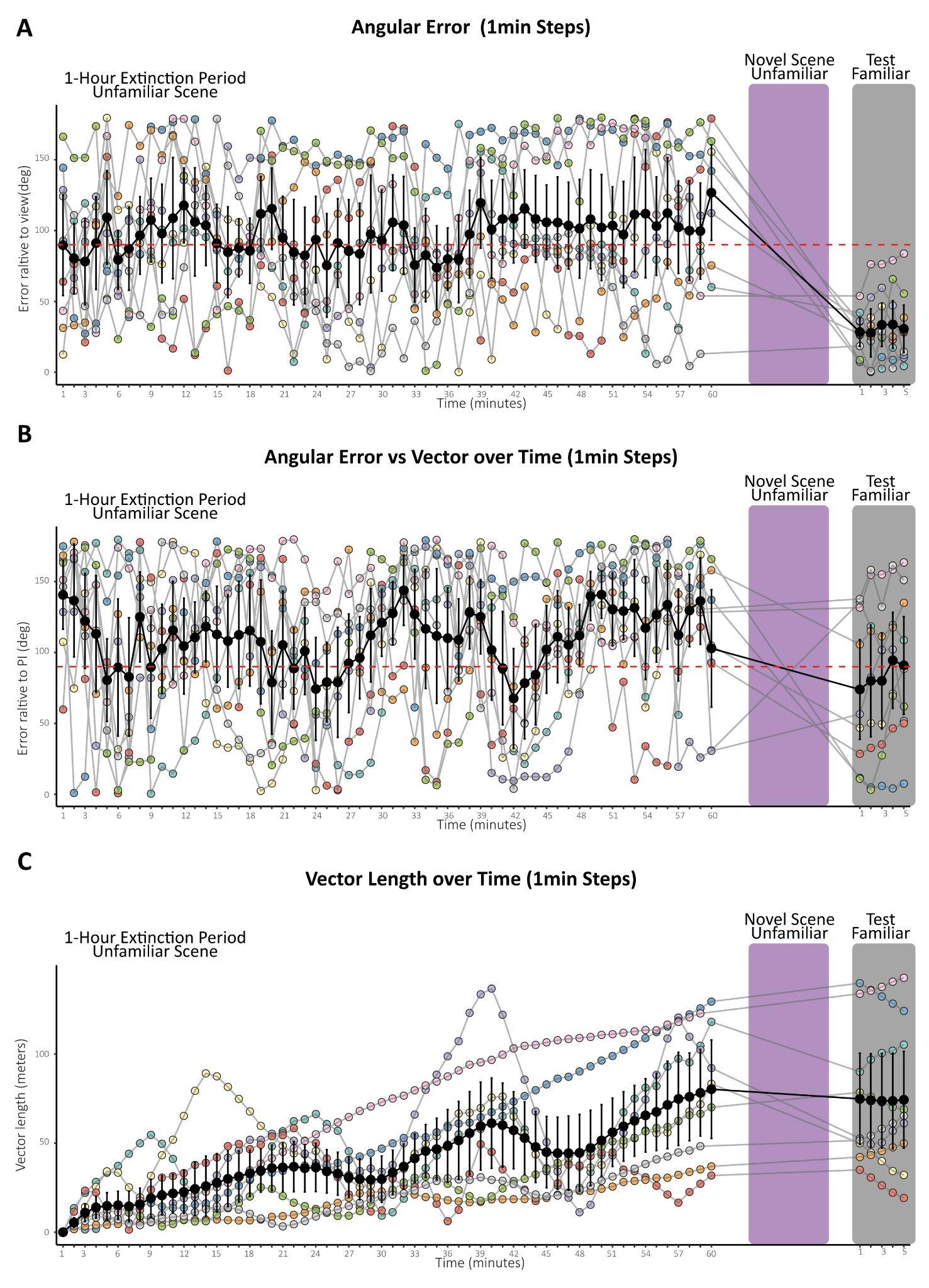

**Supplemental Figure 3. Heading error relative to nest-aligned views and the Path Integrator (PI) in 1-minute increments during a 1-hour exposure to an unfamiliar site. (A)** Angular Error vs Nest Compass Direction. Foragers showed no orientation to the nest compass at the unfamiliar sites (A and B), but immediately oriented to the nest-aligned view upon transfer to the familiar route (grey). This confirms sustained motivation. **(B)** PI Orientation. Foragers exhibited no consistent orientation toward their accumulated PI vector across any phase of the experiment. **(C)** Vector Length over time. PI vector length increased steadily throughout the 1-hour trackball session. This confirms that the lack of orientation was not due to foragers "running down" their vector, but rather a lack of reliance on the PI in an unfamiliar context.

Black line, mean. Bars, SE. The red-dotted line denotes the orientation threshold.

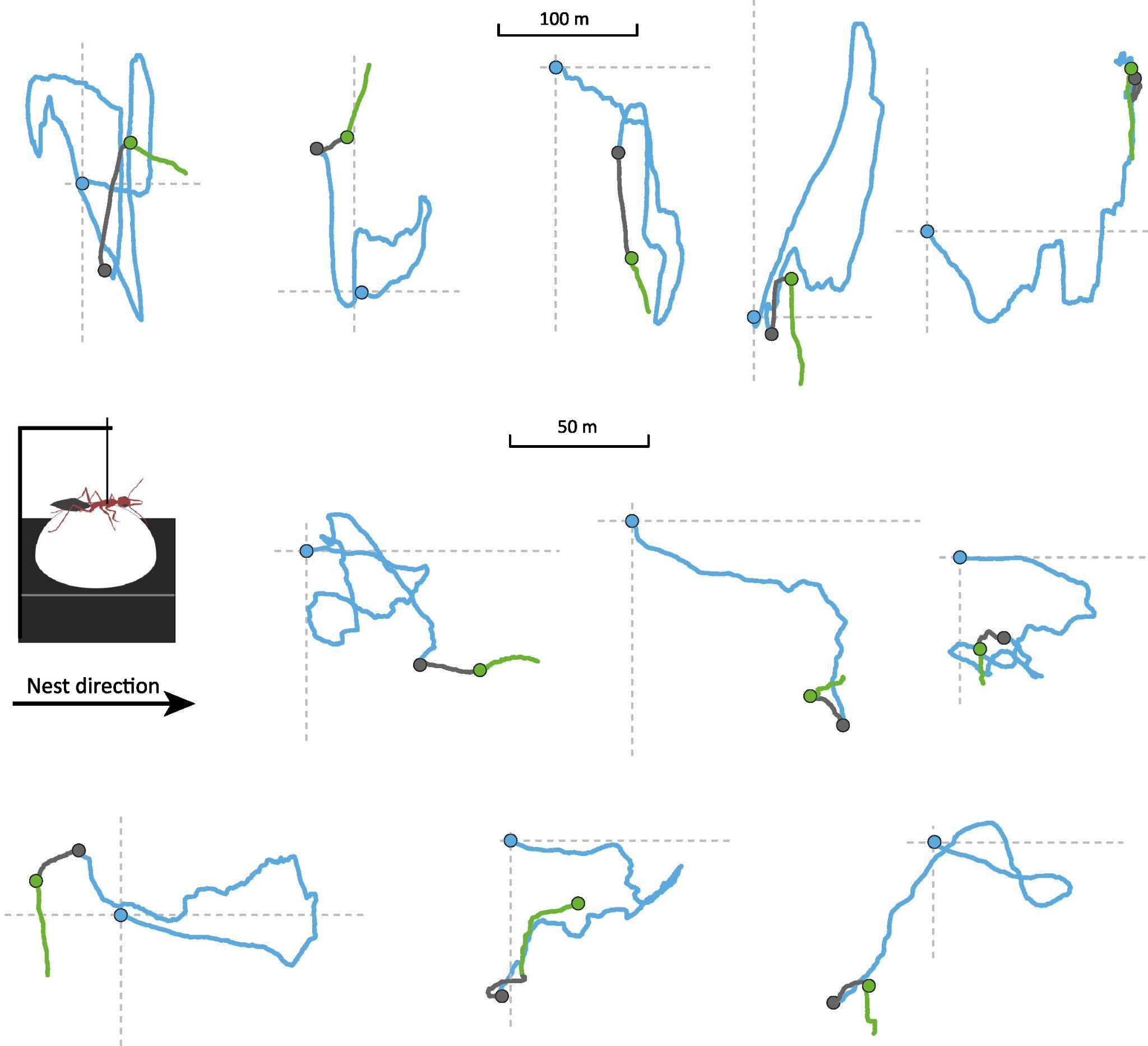

**Supplemental Figure 4. Representative homing paths of tethered individuals in the 1-hour extinction condition at an unfamiliar site.** Blue paths indicate initial nest-alignment during the first ~20 minutes of homing following tethering (blue circle) and the progressive decline of nest-alignment. After one hour, foragers were transported to a distant unfamiliar site (grey dot); grey paths show movement during this 5-minute novel scene exposure period. Finally, individuals were returned to the original extinction site (green dot); green paths show the subsequent 5-minute re-exposure period to the original scene.
